# Temperature is a dominant driver of distinct annual seasonality of leaf litter production of equatorial tropical rain forests

**DOI:** 10.1101/454058

**Authors:** Kanehiro Kitayama, Masayuki Ushio, Shin-ichiro Aiba

## Abstract

1. Intra-annual periodicity of canopy photosynthetic activity and leaf development has been documented in seasonal and weakly-seasonal tropical forests in the Amazon and elsewhere. However, vegetative periodicity such as leaf flush and fall in apparently “aseasonal” equatorial tropical forests has not been well documented. Moreover, causal drivers of the vegetative periodicity in those forests have not been identified largely because of the difficulty in performing manipulative experiments targeting whole forest ecosystem dynamics.
2. Here we show a distinct annual seasonality in canopy dynamics using a Fourier analysis with a statistical significance test on the long-term, fortnightly monitored dataset of leaf litterfall in nine evergreen tropical rain forests on Mount Kinabalu, Borneo. Statistically significant annual periodicity occurs across altitudes and soil types in all years irrespective of the year-to-year climatic variability, suggesting that fluctuations in regional climate rather than local micro-climatic, edaphic and/or biotic conditions cause the precise 1-year periodicity.
3. We examine climatic factors that have causative effects on the distinct 1-year periodicity using the spectrum convergent cross mapping that we developed in the present study to distinguish causal relationships from seasonality-driven synchronization. According to the analysis, we find that mean daily air temperature is most strongly, causatively related to the 1-year periodicity of leaf litterfall. However, knowledge on ecophysiolocial and molecular mechanisms underlying temperature-control of tropical tree growth is limited and further studies are required to understand the detailed mechanisms.
4. (*Synthesis*) We suggest that intra-annual temperature changes in association with the movement of the intertropical convergence zone cause the distinct annual vegetative periodicity. Because vegetative periodicity can be transmitted to the dynamics of higher trophic levels through a trophic cascade, interactions between vegetative periodicity and daily air temperature, not rainfall, would more strongly cause changes in the dynamics of equatorial tropical rain forests. Our results show that clear vegetative periodicity (i.e., annual seasonality) can be found in equatorial tropical rain forests under diverse local environments, and that air temperature is a more important factor than the other climate variables in the climate-forest ecosystem interactions.

## 1 INTRODUCTION

One of the most well-defined characteristics of the equatorial tropical rain-forest climate is the lack of clear annual changes in mean monthly temperature, while monthly rainfall always exceeds monthly potential evapotranspiration (Whitmore 1975). This hot and humid climate throughout the year without marked seasonality supports vegetative growth year round in tropical rain forests, leading to high primary productivity (Whitmore 1975).

In spite of the year-round constant climate, tropical trees are known to show annual/supra-annual reproductive periodicity. For example, tropical trees show synchronous flowering and fruiting in the Neotropics (Borchert et al. 2005) and Southeast Asia (Sakai 2002, Sakai et al. 2006). Seasonal radiation change has been suggested to trigger reproductive synchrony as an annual event in the Amazon and elsewhere (Schaik et al. 1993, Wright and Calderón 2018). On the other hand, irregular El-Niño induced droughts (and sometimes, low air temperature) are suggested to trigger supra-annual flowering and fruiting in Southeast Asian tropical rain forests, the phenomenon known as masting or general flowering (Sakai 2002, Sakai et al. 2006, Cannon et al. 2007, Kobayashi et al. 2013, Chen et al. 2017, Ushio et al. 2020). Several ultimate mechanisms have been proposed for explaining masting. Massive seed crops are considered advantageous in satiating seed predators in a masting event (e.g., Janzen 1970, Kelly 1994, Kelly and Sork 2002). Synchronous reproduction is also believed to be favourable for promoting outcrossing among sparsely distributed trees within a species in tropical rain forests because the density of conspecific trees is extremely low there, reflecting the high tree-species diversity (Sakai et al. 1999). Thus, phenological changes in reproduction in such “aseasonal” rain forests have evolutionary significance through satiating seed predators and/or enhancing genetic exchanges.

Distinct seasonality in vegetative activities such as canopy photosynthetic activity, and leaf flush and fall is also known in humid tropical rain forests in the Amazon (Restrepo-Coupe et al. 2013, Borchert et al. 2015, Wu et al. 2016). Such vegetative changes must have significant ecosystem consequences because changes in photosynthetic activity and synchronous leaf flush and fall influence water fluxes via changing transpiration (Wu et al. 2016), biological interactions in grazing food-chains (Schaik et al. 1993) and detritus food-chains, respectively (Lodge et al. 1994). Insolation and photoperiodicity and their influences on temperature might drive the annual seasonality according to correlation analyses (Rivera et al. 2002, Restrepo-Coupe et al. 2013, Borchert et al. 2015, Wagner et al. 2017). However, other studies have suggested that leaf-level phenology, not seasonality of climate, might be a driving factor (Wu et al. 2016), and thus causal factors of the annual seasonality have still been controversial. A large-scale manipulative experiment, an effective strategy to prove causal relationship, was conducted in a Panamanian tropical forest, where desiccation had been identified as a stress factor, to investigate the response of litterfall to ameliorated soil desiccation with artificial water supplies in two dry seasons (Wright and Calderón 2018). Nonetheless, if candidate causal drivers are multiple and do not involve stress factors, a manipulative experiment is usually impractical for gigantic ecosystems such as tropical rain forests and researchers rely on correlation analyses to infer causal drivers of the annual seasonality. In addition, although the annual seasonality has been often reported in Amazonian forests, descriptions of annual vegetative periodicity are still limited in equatorial forests in Southeast Asia (but see Nakagawa et al. 2019). Therefore, whether the annual vegetative periodicity is widespread in equatorial tropical forests and how the vegetative periodicity is driven are still unknown.

In the present study, we examine vegetative periodicity and its causal factor in equatorial tropical rainforests in Borneo using a long-term litterfall and meteorological monitoring data. Our research sites are on an altitudinal gradient on Mount Kinabalu (from 700 to 3100 m) and cover a wide range of climatic and edaphic conditions, which enables evaluation of vegetative periodicity under various ecological conditions. As will be explained later, our sites consist of a matrix of four altitudes and two soil types (soils derived from sedimentary vs. ultrabasic rocks). Sedimentary soils are relatively richer in soil phosphorus availability than ultrabasic soils (Aiba and Kitayama 1999, Kitayama and Aiba 2002). We can evaluate the interactions of altitudes and phosphorus availability on vegetative periodicity using this setting. Above-ground net primary productivity was lower on ultrabasic than on sedimentary soils at the same altitude, and sharply decreased with increasing altitude on both soils (Kitayama and Aiba 2002). Trees show more resource-conservative traits on P-deficient, low-productivity forests across our study sites (Tsujii et al. 2020). Particularly, we were interested to know if/how resource-conservation traits of trees influence vegetative periodicity. Resource conservation mechanisms would favour a longer periodicity even climatic cues are the same because timing and amount of resource allocation would determine the patterns of vegetative periodicity. Thus, we expected variable periodicities depending on productivity across our sites in association with altitudes and soils. In the present study, first, we show that, using a Fourier analysis with a significance test (Bush et al. 2017), leaf litterfall patterns show a distinct annual seasonality in the Bornean tropical rain forests. Second, we apply a newly developed causal analysis that can distinguish causal relationships from seasonality-driven synchronization to the long-term litterfall and meteorological monitoring data and show that air temperature, not rainfall, is a dominant driver of the vegetative periodicity in the equatorial tropical rain forests.

## 2 MATERIALS AND METHODS

### 2.1 Study sites and litter collection

The study sites are located on Mount Kinabalu (4095 m, 6 5 N, 116 33 E), Borneo, which is a non-volcanic mountain consisting of Tertiary sedimentary rocks of sandstone and/or mudstone below 3000 m and of granitic rock (also of Tertiary origin) above 3000 m. Ultrabasic rocks (Mesozoic serpentinite) also occur as mosaics below 3100 m. A pair of forests were selected on two contrasting geological substrates (one derived from sedimentary rock and the other derived from ultrabasic rock) each at 700, 1700, 2700 and 3100 m on the southern slope. The 3100-m “sedimentary” site is actually underlain by granitic rocks. In this paper, sedimentary/granite site is hereafter referred to as “sedimentary site”. These forests consist of a matrix of two soil types and four altitudes. In addition, we added a ninth forest at 1700 m on Quaternary sediments originated from alluvium/colluvium deposits. The concentrations of both total phosphorus and soluble phosphorus are greater in sedimentary soils than in ultrabasic soils at the same altitude (Kitayama et al. 2000); these concentrations decrease in the order of the Quaternary site (richest phosphorus), the sedimentary site and the ultrabasic site (poorest phosphorus) at 1700 m. Forests consist of lowland tropical rain forests at 700 m, lower montane tropical rain forests at 1700 m, upper montane tropical rain forests at 2700 m, and subalpine forests at 3100 m. Dominant families are Dipterocarpaceae at 700 m and Fagaceae and Myrtaceae above 1700 m. Evergreen gymnosperms of the family Podocarpaceae are co-dominant at the 1700-m sedimentary site, as well as at the three ultrabasic sites above 1700 m. The structure and species composition of these forests were described in detail in the previous papers (Aiba and Kitayama 1999, Kitayama and Aiba 2002).

### 2.2 Litter monitoring data

Litterfall monitoring was performed from February 1996 until March 2006 or longer at all sites (Figure 1, Table S1) except for the 700-m ultrabasic site, where monitoring was abandoned after two years due to the site’s extreme remoteness. Detailed descriptions of data collection are given in a previous paper (Kitayama & Aiba, 2002). Briefly, a total of 20 litter traps at two lower altitudes, and a total of 10 traps at two higher altitudes were placed 1 m above the ground at 10 m intervals along 2 transects at each site. Traps were made of 1 mm mesh with an opening of 0.5 m^2^ area, and were renewed annually. Field assistants collected trapped fine litter from each trap at two-week intervals without interruption from all sites. Collected litter was immediately oven-dried at 70°C for at least 3 days, sorted by trap into six fractions (leaves, reproductive organs including fruits and flowers, twigs ≥ 2 cm girth, epiphytes, palms and bamboos, and dust), and weighed to determine dry weight. We applied time-series analyses to each fraction in each forest.

**Figure 1.**
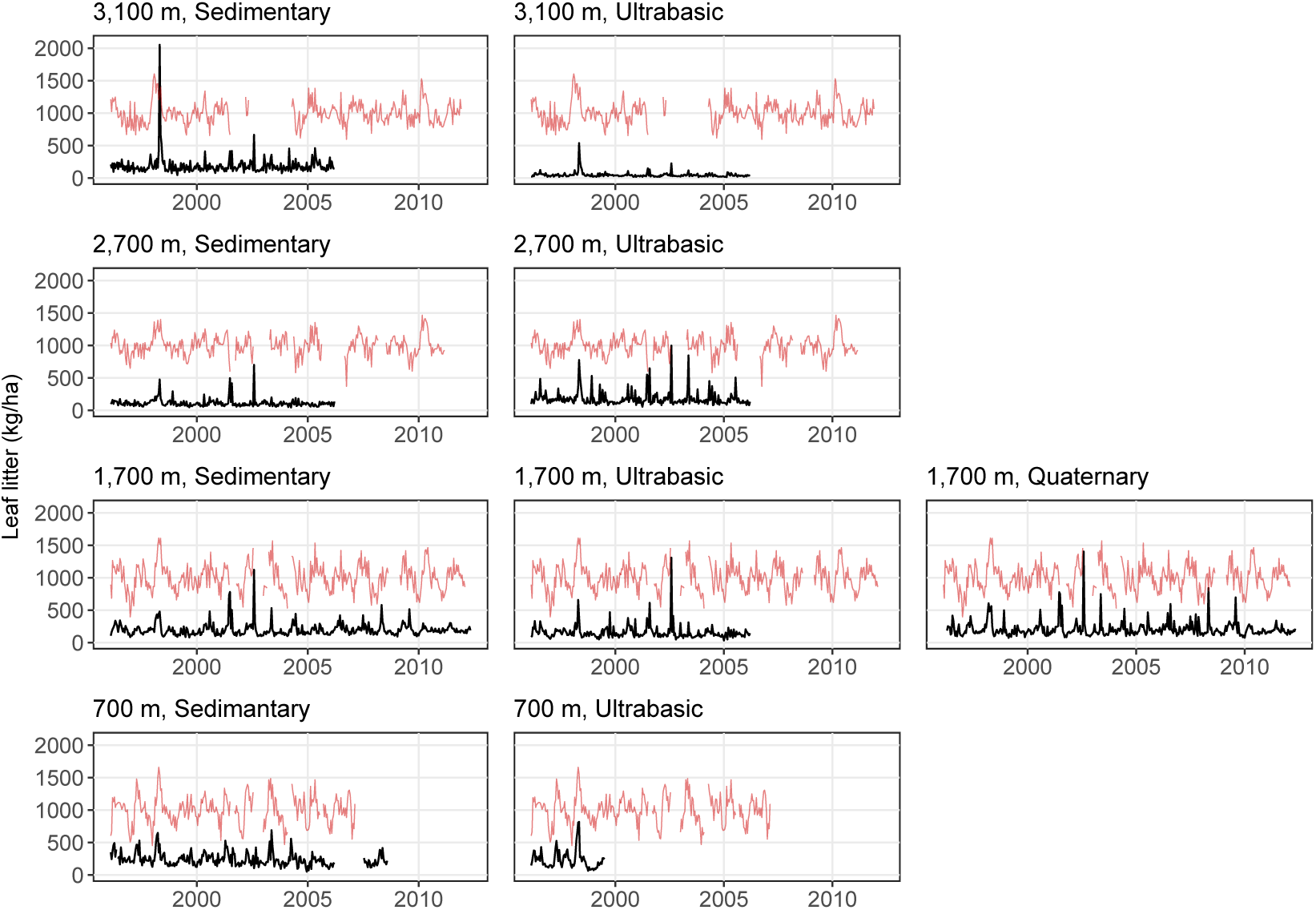
Time series of leaf litter production in tropical rain forests on Mt. Kinabalu. Black lines indicate leaf litter production, and red lines indicate mean daily air temperature corrected by a general additive model (only patterns are shown). The values on the -axis are for leaf litter production.

Because we performed the litter collections at two-week intervals, there are 26.07 sampling events per year, or 26.14 sampling events per year in leap years. In order to make our analysis simple and straightforward, we converted the original data to time series with 24 time points per year throughout the whole monitoring period (i.e., constant time points per year for the whole monitoring period). Briefly, each month was divided into two time periods (e.g., early January and late January), and data points were assigned into either time period in each month. Most time periods include only one sampling event. If a time period includes two sampling events, we calculated the mean value, and it was assigned as a representative value of the time period. The converted time series had 24 time points per year throughout the monitoring period, and the subsequent analysis and interpretations of the results would be easier than those with the original time series.

### 2.3 Meteorological data

Four automated meteorological stations were established at 550, 1560, 2700 and 3270 m on the south slope. Each station was located in an open place without obstacles within at least a 10-m radius. Each station consisted of climate sensors connected to a CR10X data logger (Campbell Scientific, Logan, Utah, USA); a Vaisala HMP35C probe for air temperature and relative humidity, a 107B probe for soil temperature (10 cm), a LI-COR 190SB quantum sensor for photosynthetically active radiation (PAR) (wavelength 400-700 nm), a TE525MM tipping bucket rain gauge for rainfall, and an RM Young 03001 Wind Sentry for wind speed/direction. All readings were taken at 10-s intervals, and reduced to 30-min and 24-hour means and/or totals, and electronically stored in CR10X. Saturation deficits were calculated based on 30-min mean air temperature and instantaneous relative humidity values. Daily potential evapotranspiration was estimated following the Penman-Monteith equation (Allen et al. 1998). Some of the sensors were renewed at regular intervals; however, baseline drifts due to sensor deterioration were still inevitable for mean daily PAR and saturation deficit.

The baseline drifts due to sensor deterioration may influence the performance of time series analysis. The baseline drifts were, therefore, corrected by constructing additive models using the ‘mgcv’ package (Wood 2004) of R. In the corrections, we assumed that the baseline drifts are long-term fluctuations in the climate variables, and that the additive models with the appropriate number of knots (*k*) can represent the long-term fluctuations. The additive models successfully represented the baseline drifts in the climate time series (correction results are available at https://github.com/ong8181/kinabalu-spectrum-CCM). Also, removing long-term trends made time series more stationary, which may allow better detection of dominant cycles and causality between time series. The residuals of the additive models do not include the long-term trends and were used as corrected climate time series for the subsequent time series analysis.

### 2.4 Fourier analysis

We performed a Fourier analysis to identify dominant cycles of meteorological and litter time series in the tropical forests. Fourier is a spectral analysis used to decompose a time series into sine waves of different frequencies. Because Fourier analysis requires consecutive time series (i.e., missing values are not allowed), we selected a longest consecutive time series (Table S1). As meteorological time series sometimes contain missing values, a maximum of six consecutive missing values were interpolated by using simple linear interpolation (the maximum number of consecutive missing value was determined by taking the balance between the length of time series and artifacts introduced by the linear interpolation). For litter time series, there were no missing values.

Bush et al. (2017) proposed a practical procedure to detect a dominant cycle from a given time series, and we generally followed that method. First, the raw spectral estimates were smoothed using the modified Daniell kernel. We adjusted the width of the Daniell kernel so that a bandwidth become approximately 0.1, which was found to give sufficient resolution to identify a dominant peak (Bush et al. 2017). The significance of the dominant peak was tested by calculating 95% confidence intervals, assuming that spectral estimates approximate a chi-square distribution (Bloomfield 2000). We rely on the null continuum of the spectrum (i.e., the null spectrum) as a null hypothesis rather than the average spectrum (i.e., white noise spectrum) because the null spectrum showed fewer false positive results (Bush et al. 2017). See Tables S2 and S3 for the summary of Fourier analysis. In addition to the Fourier analysis, we performed the wavelet analysis to supplement the Fourier analysis and visualize how the seasonality strength changes over time. Morlet wavelet was used as a mother wavelet, and this analysis was performed using ‘WaveletComp’ package of R (Röesch and Schmidbauer 2018).

### 2.5 Time series of seasonality and the framework of spectrum convergent cross mapping (spectrum CCM)

As will be shown, our analysis showed that the dominant cycles of meteorological and litter (especially, leaf litter) time series were 12 months (i.e., annual cycle) at many sites. Synchronized seasonality often makes detection of causality extremely difficult. Indeed, previous studies showed that synchronization driven by seasonality can lead to misidentification of causality (i.e., false-positive detection of causality) (Deyle et al. 2016a, Cobey et al. 2016, Ushio et al. 2018). Several methods have been proposed to overcome this limitation (Deyle et al. 2016a, Ushio et al. 2018), but the issue is still unresolved. More importantly, even if a causal variable is identified, the variable may not necessarily be a driver of “seasonality”. For example, if a causal variable has influences only during a certain period of the year, then the variable only slightly contributes to the whole seasonality of an effect variable. Such variable is a causal factor, but not a driver of seasonality of a response variable.

To overcome this problem, here we introduce spectrum convergent cross mapping (spectrum CCM) as a general approach to detect a driver of seasonality among highly synchronized time series. The idea is straight-forward: if a time series is a driver of seasonality of another time series, then the seasonality of the causal time series drives the seasonality of the effect time series. Strength of seasonality, which can be quantified as the power of an annual cycle (i.e., the amplitude of seasonality), can also be a time series if we sequentially calculate using a moving window method, and thus CCM, a causality test, allows to detect causality among the powers of seasonality (for CCM, see the following section).

Time series of seasonality were generated as foellows (Figure S1): (1) A 3-year window was set in an original time series (the size of the window, 3-year, was determined by taking the balance between the length of time series and the reliability of the seasonality quantification), (2) Fourier analysis was performed for the 3-year time series, (3) the power of annual cycle (i.e., seasonality) was extracted, (4) steps 1–3 were sequentially performed for the whole time series, and (5) the obtained time series were then first-differenced to make the time series stationary. Briefly, when the total length (time point) of time series *X* with time interval at 2-weeks is *N*, the ith 3-year window is described as follows,

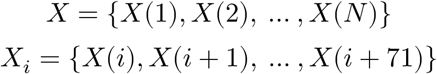

where *X*(*i*) indicates the *i*th element of the time series and *X*_*i*_ should contain 24 × 3 = 72 elements corresponding to the 3-year term (Figure S1a). In total, we obtain *N*–71 of *X*_*i*_ and Fourier analysis is applied to each *X*_*i*_, resulting in *N*–71 powers of the 1-year periodicity (Figure S1b). Let the power of the 1-year periodicity of *X*_*i*_ be *S*(*i*) and time series of *S*(*i*) be *S*, and then the first-differenced *S* is described as follows (Figure S1c, d),

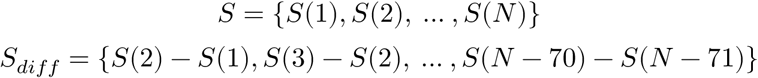

*S*_*diff*_ is normalized to zero mean and unit variance and then used as an input for CCM explained in the next subsection.

### 2.6 Spectrum convergent cross mapping (CCM)

Causalities among the first-differenced time series of seasonality were tested with convergent cross mapping (CCM) (Sugihara et al. 2012). CCM is based on Takens’ theorem for nonlinear dynamical systems (Takens 1981). Consider a multi-variable dynamical system, in which only some of variables are observable. Takens’ theorem (Takens 1981), with several extensions (Deyle and Sugihara 2011), proved that it is possible to represent the system dynamics in a state space by substituting time lags of the observable variables for the unknown variables. This gives a time-delayed coordinate representation (or embedding) of the system trajectories, and this operation is referred to as state space reconstruction (SSR). For example, consider a multi-variable system including a variable *X*. Even if other variables involved in the system are unobserved, the time-delayed embedding of *X*, that is {*X*(*t*), *X*(*t*−1), *X*(*t*−2), …, *X*(*t*− (*E* − 1))} where E is the embedding dimension, represent the whole dynamics of the multi-variable system. Nonlinear analytical tools have been developed based on SSR of empirical dynamics, and this framework is called empirical dynamic modeling (EDM) (Sugihara et al. 2012, Ye et al. 2015). Recent studies demonstrated that EDM is a useful tool to analyze nonlinear dynamics in natural ecosystems (Ye et al. 2015, Deyle et al. 2016b, Chang et al. 2017, Ushio et al. 2018, Ushio 2020).

An important consequence of the SSR theorems is that if two variables are part of the same dynamical system, then the reconstructed state spaces of the two variables will represent topologically the same attractor. Therefore, it is possible to predict the current state of a variable using time lags of another variable. We can look for the signature of a causal variable in the time series of an effect variable by testing whether there is a correspondence between their reconstructed state spaces (i.e., cross mapping). Cross-map skill is evaluated by a correlation coefficient (*ρ*) between observed and predicted values by cross mapping. In addition, the cross-map skill will increase as the number of points in the reconstructed state space increases if two variables are causally related (i.e., convergence). In this study, cross mapping from one variable to another was performed using simplex projection (Sugihara and May 1990). How many time lags are taken in SSR (i.e, embedding dimension) is determined by simplex projection using mean absolute error as an index of forecasting skill. More detailed algorithms about simplex projection and cross mapping can be found in previous studies (Sugihara et al. 2012, Ye et al. 2015, Chang et al. 2017).

The significance of CCM is judged by comparing cross-map skill and convergence of Fourier surrogates and original time series. More specifically, first, 1000 surrogate time series for one original time series are generated. Fourier surrogates were generated by randomizing the phases of a Fourier transform using `rEDM::make_surrogate_ebisuzaki()` function implemented in rEDM package of R (Ye et al. 2018). Second, cross-map skill and convergence (i.e., the difference between cross-map skills (*ρ*) at the minimum and maximum library lengths) are calculated for 1000 surrogate time series and the original time series. If the number of surrogates that show a higher *ρ* as well as a higher convergence is less than 50 (i.e., 5% of the surrogates), the cross mapping is judged as significant. Hereafter, the number of surrogates that show a higher ρ as well as a higher convergence is referred to as “joint P-value”. We evaluated the performance of the spectrum CCM (i.e., inputs are time series of the strength of seasonality) and normal CCM (i.e., inputs are raw time series) using simulation data. The results showed that the spectrum CCM indeed distinguishes the true driver of seasonality from synchronized, non-driver variables (Supplementary Information, Figures S2–3).

### 2.7 Computation

CCM was performed using the ‘rEDM’ package (version 0.7.1) (Ye et al. 2018), and all statistical analyses were performed in the free statistical environment R v3.4.4 (R Core Team 2018).

## 3 RESULTS

### 3.1 Seasonality of litterfall and meteorological data

Fourier analysis detected the presence of significant 12 mo (or 11.8 mo) length of periodicity in leaf litterfall in all forests except for the 2700-m sedimentary and the 3100-m sedimentary sites, where the dominant cycle was 6.3 and 24 mo, respectively (Figures 2a–d, S4–S7, and Table S2). For the 700-m ultrabasic site, the dominant periodicity was 12 mo, but it was not significant due to insufficient length of the time series, and thus we omitted the 700-m ultrabasic site from the subsequent analyses (Figure 2d). Exact (or nearly exact) and significant 12-mo periodicity occurred irrespective of altitude and soil type in six out of the eight forests, suggesting that the periodicity is not influenced by local climatic, edaphic conditions, and/or their interactions. By contrast, significant dominant cycles of reproductive-organ litter (flowers and fruits combined) range from 15 mo to more than 48 mo, and periodicity was not consistent across altitudes and soil types (Figure 2a–d).

**Figure 2.**
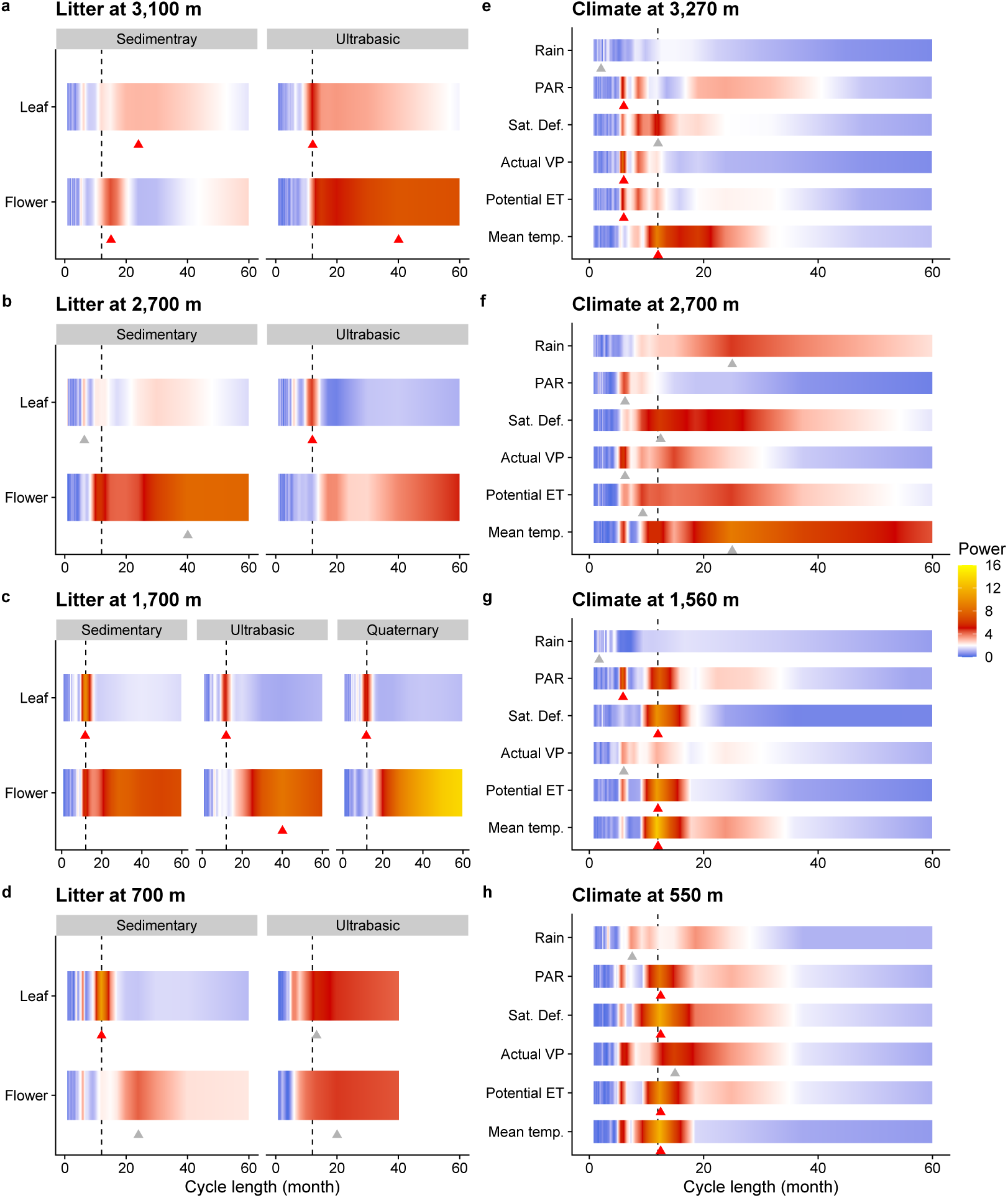
Significance of the dominant cycle and spectrum of each time series. Spectrum of litter time series collected (**a**) at sedimentary and ultrabasic sites at 3100 m, (**b**) at sedimentary and ultrabasic sites at 2700 m, (**c**) at sedimentary, ultrabasic and Quaternary sites at 1700 m and (**d**) at sedimentary and ultrabasic sites at 700 m. Note that litter monitoring was abandoned at the ultrabasic site at 700 m after two years, and therefore the time series length is much shorter than those of the other sites. Spectrum of climate variables measured (**e**) at 3270 m, (**f**) 2700 m, (**g**) 1560 m, and (**h**) 550 m (corresponding to 3100 m, 2700 m, 1700 m and 700 m forest site, respectively). Mean temp., Potential ET, Actual VP, Sat. Def., PAR and Rain indicate mean daily air temperature, potential evapotranspiration, actual vapor pressure, saturation deficit, photosynthetically active radiation, and daily rainfall, respectively. Dashed vertical line in each panel indicates the 1-year periodicity. Red and gray triangles indicate significant dominant periodicity and insignificant dominant periodicity at P < 0.05, respectively (see more details in Figure S7). Dominant periodicity longer than 60 mo is not shown here, but presented in Tables S2 and S3. Gradient colours indicate the strength of the power.

As for the meteorological variables, a significant 12-mo periodicity was found for mean daily air temperature at 550, 1560, and 3270 m (Figure 2e–h, Table S3), for saturation deficit at 550 and 1560 m, for daily potential evapotranspiration at 550 and 1560 m (Figure 2g–h), and for mean daily PAR at 550 m only (Figure 2h). Daily rainfall did not show a 12-mo periodicity at any altitude. The time series were also analyzed by the wavelet analysis, of which results were generally consistent with those of the Fourier analysis (Figures S8–S10). For example, for leaf litter production, strong 1-year periodicity was detected at seven out of the nine study sites (Figure S8). On the other hand, no clear periodicity was detected for flower litter production (Figure S9). Mean daily air temperature showed relatively strong one-year periodicity for all sites (Figure S10).

### 3.1 Causal drivers of the litterfall seasonality

The spectrum CCM analyses of the litterfall and meteorological datasets identify meteorological factors that cause the precise 1-year (or 12-mo) periodicity of leaf litterfall in the six forests out of eight forests we examined (note that the 700 m ultrabasic site was not included in the analysis because of the short time series). Mean daily air temperature is causally related to the annual periodicity of leaf litterfall across all forests except for the 1700-m sedimentary forest (Figure 3). Time lags between responses (leaf litterfall dynamics) and signals (mean daily air temperature) range from 1 month (leaf fall lags behind mean daily air temperature by 1 month) at 700 m to 7.5 months or longer at higher altitudes (Figure 3). The 1-year periodicity of leaf litterfall in the 700-m sedimentary forest is causally related to potential evapotranspiration, saturation deficit and PAR in addition to mean daily air temperature. In addition, mean daily air temperature shows the highest cross-map skill (*ρ*) within each forest, suggesting that mean daily air temperature is a leading factor that drives the 1-year periodicity of leaf litterfall (Figure 3).

**Figure 3.**
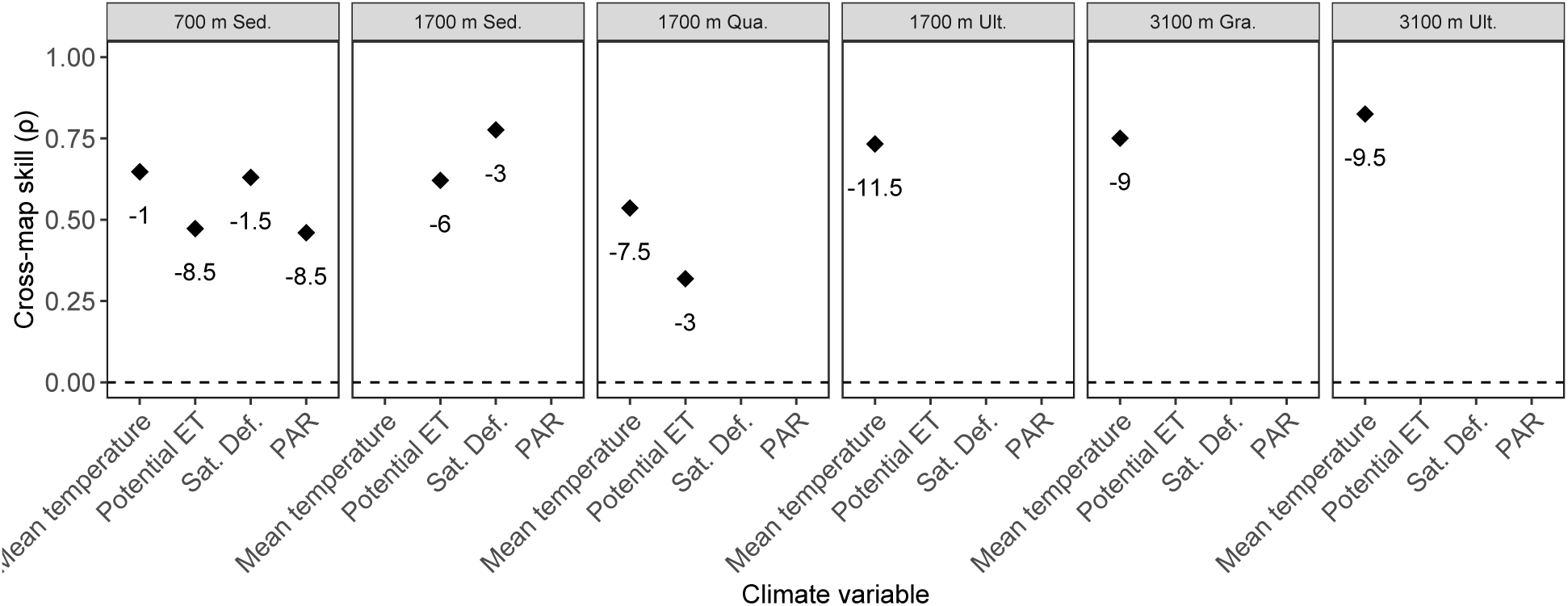
Results of spectrum convergent cross mapping (spectrum CCM). Points indicate cross-map skills (ρ; correlation coefficient between observed values predicted by the cross-mapping). Only significant (P < 0.05) cross-map skills are shown. The number below each point indicates time-lag (months) of the significant cross-mapping. Labels on the x-axis indicate climate variables. The label on the top of each panel indicates the forest site.

## 4 DISCUSSION

In the present study, first, we demonstrate that leaf litterfall patterns show a distinct annual seasonality in the Bornean tropical rain forests under a wide range of climatic and edaphic conditions (Figure 2). Second, we developed a new causal analytical framework to distinguish causal relationships from seasonality-driven synchronization, and show that air temperature is a dominant driver of the annual periodicity in the equatorial tropical rain forests (Figure 3).

The occurrence of the distinct annual leaf-litterfall seasonality across wide environmental conditions did not support our hypothesis. We originally hypothesized that trees with conservative traits in less productive forests would show a longer vegetative periodicity. Although the leaf litter production at 3100-m sedimentary site demonstrated a 24-mo periodicity, all nutrient-impoverished forests on ultrabasic soils demonstrated a precise 1-year periodicity. This suggests that the adaptive significance of vegetative periodicity rather overrides the resource conservation mechanism in these forests and they respond to a common climatic cue.

While we found the generally consistent intra-annual periodicity of leaf litterfall and meteorological variables across the elevation gradient on Mount Kinabalu, the pattern is less clear in the two 2700 m sites (Figure 2b, f). The elevation of 2700 m corresponds to the middle of the cloud zone where thick mists envelop the forests throughout a year due to daily convective uplifts of warm air (Kitayama and Aiba 2002). Due to the nearly persistent air condensation, the periodicity of air temperature and moisture as well as radiation should be obscured at this elevation.

The precise 1-year periodicity of mean daily air temperature found at all altitudes except for 2700 m is associated with the motion of the sun following the ecliptic. However, mean monthly air temperatures peak once in May or twice in May and November in our sites (Figure S6). This suggests that the yearly course of air temperature is most likely influenced by the movement of the intertropical convergence zone (ITCZ), which crosses north Borneo twice in May and November (mean migration patterns in 17 years between 1971 and 1987; Waliser and Gautier, 1993) rather than solar cycle of surface heating. The annual cycle of the ITCZ in the western Pacific lags approximately one month behind the solar cycle of surface heating (Waliser and Gautier 1993). The influence of ITCZ as a driver of the periodicity of flowering/fruiting has also been suggested in Panama (Wright and Calderón 2018). Another possible meteorological factor causing annual periodicity is the Asiatic monsoon. However, the monsoon changes year to year in magnitude and timing, and thus it is unlikely that the monsoon is a meteorological cue for the precise 1-year periodicity (Whitmore 1975).

Mean daily air temperature as well as other meteorological variables including humidity and photosynthetically active radiation (PAR) show annual patterns in association with the ITCZ, giving rise to precise 1-year periodicity of these meteorological variables. PAR, however, does not show annual periodicity at higher altitudes (above 1560 m, Figure 2) and was not selected as a causative factor at higher altitudes, suggesting that photoperiodicity (or insolation) may not be a dominant signal for leaf dynamics at least in our study sites, unlike suggestions by previous studies (Borchert et al. 2015, Wagner et al. 2017). The difference between the previous studies (many of which were conducted in the Amazon) and our study might be due to the difference in the geographical regions of study sites. Alternatively, the previous studies might not be able to detect causal relationships because they relied on correlations. Further studies are necessary to evaluate the robustness and generality of our finding in the present study. Particularly, anomalies of the ITCZ can occur in association with the variability of the monsoon as well as the El-Niño Southern Oscillation in the western Pacific (Waliser and Gautier 1993), and how such anomalies affect the vegetative periodicity needs to be substantiated in the future.

On one hand, phenological control by mean daily air temperature must be an adaptive consequence (i.e., ultimate causes) of tropical trees to the meteorological conditions in the equatorial tropics where insolation and rainfall can be inter- and intra-annually variable due to the combined effects of synoptic (such as El-Niño) and local (such as convection) processes. Under such conditions, mean daily air temperature must be a most reliable cue for causing the annual vegetative as well as partly reproductive cycles. Phenological synchronization must be adaptive for tropical trees via avoiding predation of young leaves by satiating herbivores (e.g., Aide 1993) or promoting outcrossing of sparsely populated trees (Sakai et al. 1999).

On the other hand, however, the molecular mechanisms (i.e. proximate causes) of the temperature control of the vegetative periodicity of equatorial tropical forests are poorly understood unlike temperate herbaceous plants (Aikawa et al. 2010, Nagano et al. 2019). This is at least partly due to the much larger body size, much longer lifetime and higher diversity of tropical trees compared with those of temperate, herbaceous plants, including the model plant *Arabidopsis*, which consequently hinders the accumulation of genetic information. Although caution is needed because tropical trees may be different from Arabidopsis, molecular evidence from *Arabidopsis* systems might help to infer molecular mechanisms, and our findings are consistent with the hypothesis of decentralized circadian clocks in *Arabidopsis* (Shimizu et al. 2015). In *Arabidopsis*, circadian clocks function tissue-specifically with different meteorological signals, and the circadian clock in leaf epidermis was shown to be responsible for temperature-dependent cell elongation (Shimizu et al. 2015). Leaf litterfall is closely linked to leaf extension in a canopy, because senescent leaves on old shoots (modules consisting of leaves on a twig) are shed upon the extension of new shoots (Medway 1972, Schaik 1986, Wu et al. 2016). Thus, leaf litterfall is a proxy for leaf-cell elongation. Periodicity of leaf litterfall might be a result of the response of the circadian clock to temperature. Another study showed that a perennial herbaceous plant *Arabidopsis halleri* grown in a temperate region has more than 2800 seasonally-fluctuated genes (i.e., annual periodicity), and a growth chamber experiment showed the seasonality is largely driven by air temperature, not by light condition (Nagano et al. 2019). This study also supports our view that the air temperature is a dominant driver of the annual vegetative periodicity, but molecular studies targeting tropical trees are necessary to fill the gap between temperate herbaceous plants and tropical trees.

Although mean daily air temperature demonstrates sharp 12-mo periodicity, the mean annual amplitude of this periodicity is merely 2–3°C, with nearly 10°C daily oscillations. How plants respond to rather subtle annual temperature changes amid great daily oscillations is an intriguing question. In perennial *Arabidopsis halleri*, the expression of the flowering gene (a *FLOWRING LOCUS C* homolog) is significantly related to the accumulated temperature under a certain threshold temperature, indicating that phenological events are based on past temperature memory (Aikawa et al. 2010). The altitude-dependent time lags between response (leaf fall) and signal (temperature) in Figure 3 suggest the involvement of temperature memory based on accumulated temperature. Whether tropical trees respond to annual temperature changes by using past temperature memory needs to be answered by further ecological, genetic and statistical analyses.

In summary, our results show that the periodicity of leaf literfall in equatorial tropical rain forests is determined by mean daily air temperature, which is celestially caused by the influence of the ITCZ. Because vegetative periodicity can be transmitted to the dynamics of upper trophic levels through a trophic cascade, interactions between vegetative periodicity and daily air temperature, not rainfall, would cause changes in the dynamics of forest ecosystems. Therefore, the subtle annual periodicity in daily air temperature should be taken into account to study climate-forest ecosystem interactions in apparently “aseasonal” equatorial tropical rain forests. Whether climate change per se or increased anomalies of the El-Niño Southern Oscillation in association with climate change (Vecchi et al. 2006) in the western Pacific cause perturbation on the vegetative periodicity needs to be cautiously monitored.

## ACKNOWLEDGEMENTS

We thank Dr. Jamili Nais and Puan Rimi Repin of Sabah Parks, who have been encouraging the litter monitoring, and all other field assistants who assisted with monitoring and sample collection. This study was supported by the Global Environment Research Funds B-52 and B-11 of the Ministry of the Environment and MEXT/JSPS KAKENHI Grant Number 18KK0206 to K.K., 19H02998 to S-I.A., and the Hakubi Project in Kyoto University to M.U. We declare no conflicts of interests.

## AUTHORS’ CONTIRBUTIONS

K.K. and S.A. designed the litter monitoring and collected data; K.K. and M.U. conceived the hypothesis; M.U. invented the spectrum CCM; M.U analysed data with contribution from K.K.; K.K. and M.U. wrote the first draft; all authors were involved in interpretation, discussion and final writing.

## DATA AVAILABILITY STATEMENT

The R scripts used for the analyses and time-series of leaf and flower litter of the nine forests on Mount Kinabalu and meteorological data are available at https://github.com/ong8181/kinabalu-spectrum-CCM.

