## Supplementary Information for "Temperature is a dominant driver of distinct annual seasonality of leaf litter production of equatorial tropical rain forests"

### Contents:

- **Supplementary Methods and Results** | Performance test of spectrum CCM
- **Figure S1** | Algorithm of spectrum convergent cross mapping (spectrum CCM) and examples of model time series used for the performance test of spectrum CCM
- **Figure S2** | The results of the performance test of spectrum convergent cross mapping (spectrum CCM)
- **Figure S3** | The results of the performance test of convergent cross mapping (normal CCM)
- **Figure S4** | Monthly patterns of leaf litter production in the tropical rain forests on Mt. Kinabalu
- **Figure S5** | Time series of flower litter production in tropical rain forests on Mt. Kinabalu
- **Figure S6** | Monthly patterns of daily mean air temperature
- **Figure S7** | Confidence intervals and significance of the dominant periodicity of litter and meteorological time series
- **Figure S8** | Wavelet analysis of leaf litter production
- **Figure S9** | Wavelet analysis of flower litter production
- **Figure S10** | Wavelet analysis of daily mean air temperature
- **Table S1** | Time series length used for Fourier analysis
- **Table S2** | Summary of Fourier analysis of litter
- **Table S3** | Summary of Fourier analysis of climate variables

### Supplementary Methods and Results

#### Performance test of spectrum CCM

To evaluate the performance of the newly developed spectrum CCM (see Figure S1 for the framework of spectrum CCM), we performed a simulation test. In the simulation, we prepare four time series, including one response time series and three explaining time series with shared seasonality: 1) one time series continuously influences the response time series (time series $A$), 2) one time series influences the response time series during only a certain period of the year (time series $B$), and 3) one time series does not influence the response time series (time series $C$) (Figure S1e, f). Time series $A$, $B$, and $C$ are generated by adding a sine curve to the exponential of random values taken from a normal distribution. The amplitude of the sine curve added is defined as the strength of seasonality.

In a mathematical expression, model time series A is generated as follows:

$$R_{A,0} \sim N(0,1)$$

$$R_A = e^{R_{A,0}}$$

where $R_{A,0}$ represents random values that follow the normal distribution, and $R_A$ is the exponent of $R_{A,0}$. Seasonality is then added to make time series with shared seasonality.

$$T_A = \beta_{1,A} \times R_A + \beta_{2,A} \times Seasonality \quad \text{... (1)}$$

where $T_A$ is a generated model time series $A$, $\beta_{1,A}$ and $\beta_{2,A}$ define the relative strength of random values and seasonality, respectively, and Seasonality is defined by a sine curve ($\beta_{1,A} = 1$ in the simulation). Importantly, we add a small fluctuation in $\beta_{2,A}$ through time (i.e., coefficient of variation of $\beta_{2,A}$ is 20%; values are randomly drawn from a normal distribution, i.e., $N(1, 0.2)$, and the values are used to multiply the original $\beta_{2,A}$; full R codes are available at https://github.com/ong8181/kinabalu-spectrum-CCM) because the strength of seasonality can change in nature (this is also evident in our results). Model time series $B$ and $C$ are generated in the same way.

The response time series, $T_R$, is generated by combining a logistic map and influences time series $A$ ($T_A$) and $B$ ($T_B$).

$$T_{R,0}(t+1) = T_{R,0}(t) \times (3.8 - 3.8 \times T_{R,0}(t))$$

$$T_R(t+1) = a \times T_{R,0}(t+1) + b \times (T_A(t) - \Theta(t) \times T_B(t)) \quad \text{... (2)}$$

where $T_{R,0}$ is the internal dynamics that is not influenced by external factors, and TR represents observed values that include influences from external factors (e.g., climate factors). As shown in Eqn. (2), $T_A$ and $T_B$ have causal influences on $T_R$. As a is always one in the simulation, the relative influences of $T_A$ and $T_B$ are determined by $b$ ($b = 2$ in the simulation). The period when $T_B$ has an influence on $T_R$ is determined by $\Theta(t)$, where $\Theta(t)$ denotes the Heaviside function ($\Theta(t)$ is 0 when $t$ is the first three-fourths of a year, and is 1 during the last one-fourth of a year). Therefore, $T_B$ has causal influences on $T_R$ only during the last one fourth of a year (i.e., periodic influence). On the other hand, $T_A$ always has causal influences on $T_R$ and thus is regarded as a major driver of the seasonality of $T_R$. $T_C$ has no influence on TR.

For the performance test, we generated model time series with different seasonality strengths (i.e., $\beta_{2,i}$ where $i$ is either of $A$, $B$, or $C$; a relative amplitude of sine curve compared with non-seasonal variation) and different observation errors. Observation errors were values randomly drawn from a normal distribution as follows: $\epsilon \sim N(0, ErrorRate \times SD(T_i))$, where $SD(T_i)$

indicates the standard deviation of time series $i$ ($i$ is $A$, $B$, $C$ or $R$) and ErrorRate defines how large the errors are. Observation errors were added to all time series when $ErrorRate > 0$. We applied 11 seasonality strengths (i.e., $\beta_{2,i}$ is changed from 0 [no seasonality] to 2 [strong seasonality]; in R script, `seq(0, 2.0, by=0.2)`; see Figure S1 for example time series) and 11 observation errors (i.e., from $0\%$ to $50\%$ of standard deviation of the time series; in R script, `seq(0, 0.5, by=0.05)`) and the number of the four-time-series sets which includes $T_R$, $T_A$, $T_B$ an $T_C$ is 100 for each test. The interval of time series is two weeks, and the length of the time series is 12 years (288 time points).

For the model time series (i.e., the response time series and three explaining time series), we performed two tests. First, spectrum CCM was applied to the set of the four-time series, and we examined the $P$-value and detection probability (the proportion of time series that identified the driver of seasonality of the response time series). Second, normal CCM was applied to the set of the four-time series as a comparison (i.e., raw time series were used in CCM). Time-lag parameter of CCM is preliminary determined and one-step time-lag is used throughout the simulation (i.e., `tp = -1` in rEDM functions in R script; lagged CCM). In total, we performed 12,100,000 CCMs for spectrum CCM and normal CCM, respectively (100 replicates of model time series $\times$ 11 seasonality strengths $\times$ 11 observation errors $\times$ 1,000 Fourier surrogates for significance test = 12,100,000). Fourier surrogates were generated by randomizing the phases of a Fourier transform using `rEDM::make_surrogate_ebisuzaki()` function implemented in rEDM package of R. To evaluate the performance, "mean joint $P$-value" of 100 replicates and "detection probability" (= the number of replicates in which joint $P$-value < 0.05 among 100 replicates) were calculated.

Spectrum CCM correctly identified the true seasonality driver even under 20–30$\%$ of observation errors when the time series showed relatively weak to strong seasonality (Figure S2). Also, spectrum CCM does not detect time series $B$ and $C$ (i.e., periodic influencer and non-driver) as the driver of seasonality (a low possibility of the false-positive detection; Figure S2). Normal CCM always detects time series $A$ as a significant driver of seasonality (Figure S3a, d, g). However, normal CCM often detects time series $B$ (a periodic influencer) as the driver of seasonality, especially when the strength of seasonality is weak to strong (i.e., a high possibility of false-positive detection; Figure S3b, e, h). Also, the test showed that, even time series $C$ (a non-causal, random value time series) may be detected as the driver of seasonality when they are synchronized (Figure S3c, f, i). Overall, spectrum CCM is a much more conservative method for detecting the driver of seasonality when time series are synchronized due to shared seasonality. The strength of seasonality of our time series data that showed significant and 1-year periodicity is approximately in a range of relatively weak to modest seasonality (Figures 1, S1e), and thus, the application of spectrum CCM to our data enables reliable detection of a driver of seasonality.

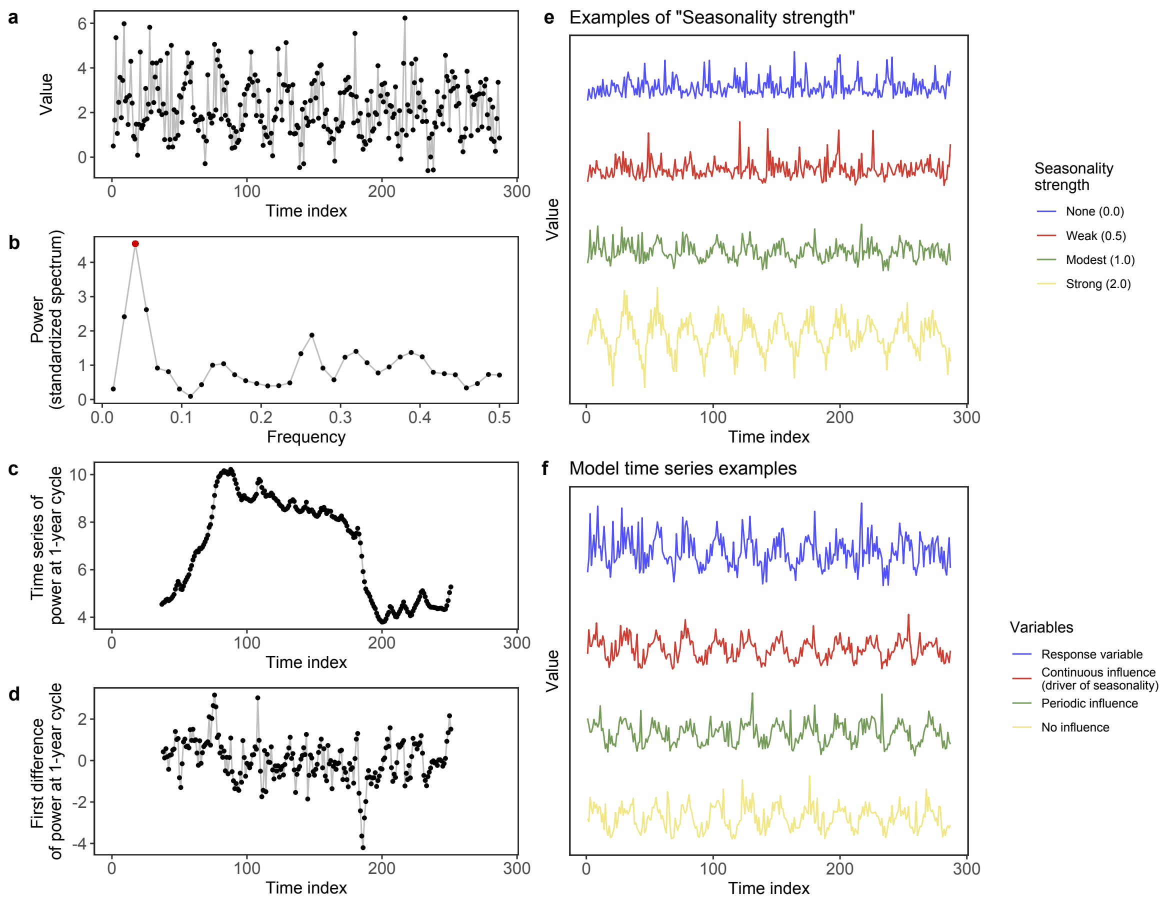

**Figure S1 | Algorithm of spectrum convergent cross mapping (spectrum CCM) and examples of model time series used for the performance test of spectrum CCM.** (**a**) Model time series that shows moderate seasonality (twelve-year time series with 2-week intervals). (**b**) Fourier analysis of the model time series, showing that the length of the dominant cycle is 12 months. Red point indicates the power of the 12-month cycle. (**c**) Time series of the power of one-year cycle calculated as follows: (1) power of one-year cycle is calculated for a three-year window, (2) we slid the window one time-step forward, (3) power of one-year cycle is again calculated for the new three-year window, and (4) steps 2-3 were repeated until the end of the time series. (**d**) First difference of the time series of the power of one-year cycle. CCM was performed for the first difference time series. (**e**) Examples of the model time series with different strengths of seasonality. Numbers in parentheses indicate the strength of seasonality (corresponding to $\beta_{2,i}$ in Eqn. (1) in Methods). (**f**) Examples of the model time series. Blue line indicates a response time series (e.g., leaf litter time series; Eqn. (2) in Methods). Red, green, and yellow lines indicate time series A, B and C (see Eqn. (1) in Methods). Time series $A$, $B$ and $C$ have continuous, periodic, and no influence on the response variable, respectively. The red time series is a true driver of the seasonality of the response variable.

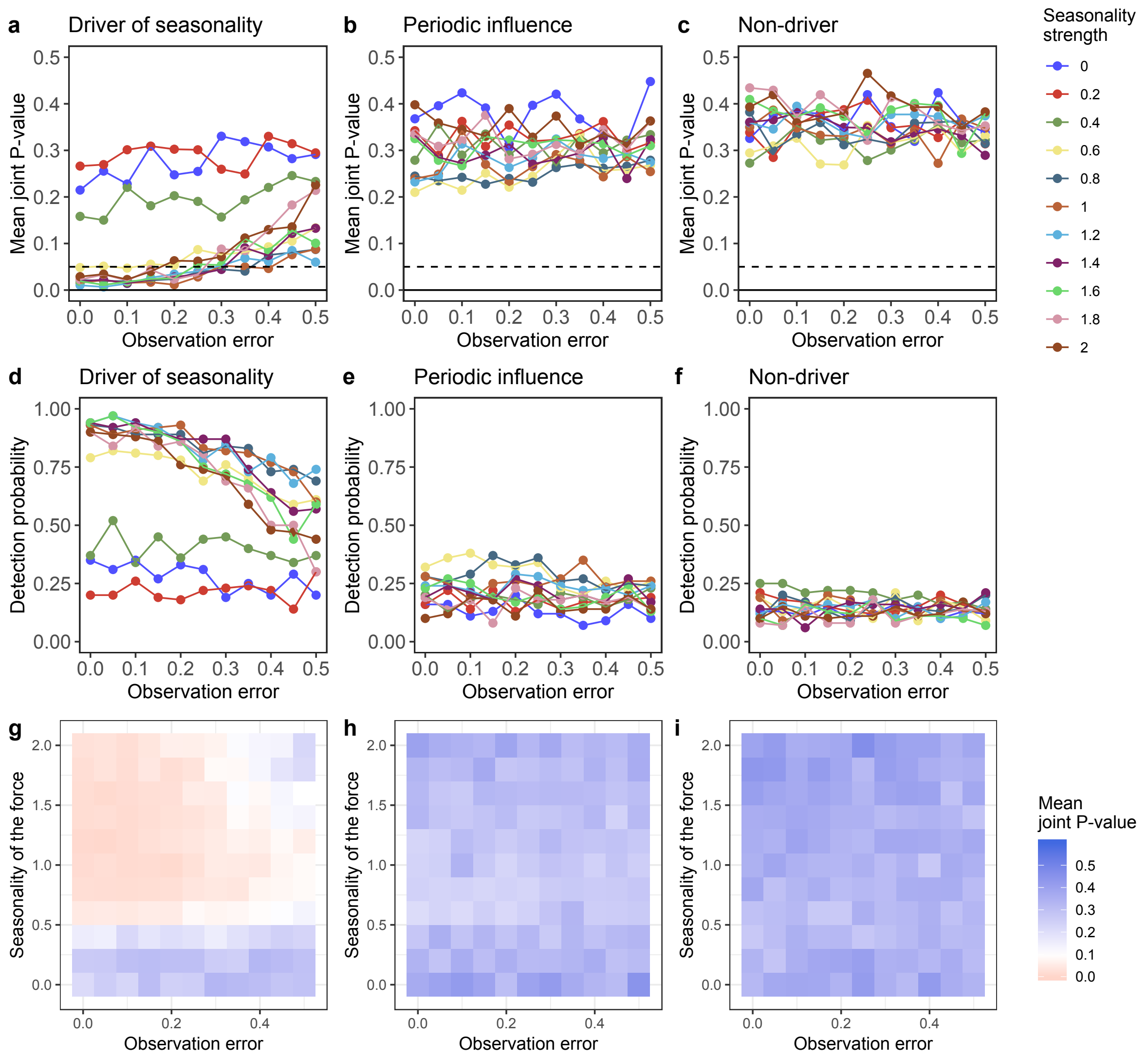

**Figure S2 | The results of the performance test of spectrum convergent cross mapping (spectrum CCM).** Seasonality strength, observation errors and mean joint $P$-values of spectrum CCM applied to time series that have (**a**) a continuous influence (the true seasonality driver), (**b**) a periodic influence, and (**c**) no influence on the target variable. Different colours indicate different strengths of seasonality. Dashed line indicates joint $P$-value = 0.05. Seasonality strength, observation errors and detection probability (as the true seasonality driver) of spectrum CCM applied to time series that have (**d**) a continuous influence (the true seasonality driver), (**e**) a periodic influence, and (**f**) no influence on the target variable. Different colours indicate different strengths of seasonality. Heat map of the seasonality strength, observation errors and mean joint $P$-values of spectrum CCM applied to time series that have (**g**) a continuous influence (the true seasonality driver), (**h**) a periodic influence, and (**i**) no influence on the target variable. Colours indicate mean joint $P$-value. If a time series has weak to strong seasonality and observation errors less than 20-30%, then spectrum CCM correctly distinguishes the true seasonality driver from non-driver (**a**–**c**), and the probability of false-positive detection is low (10–20%; **e**–**f**).

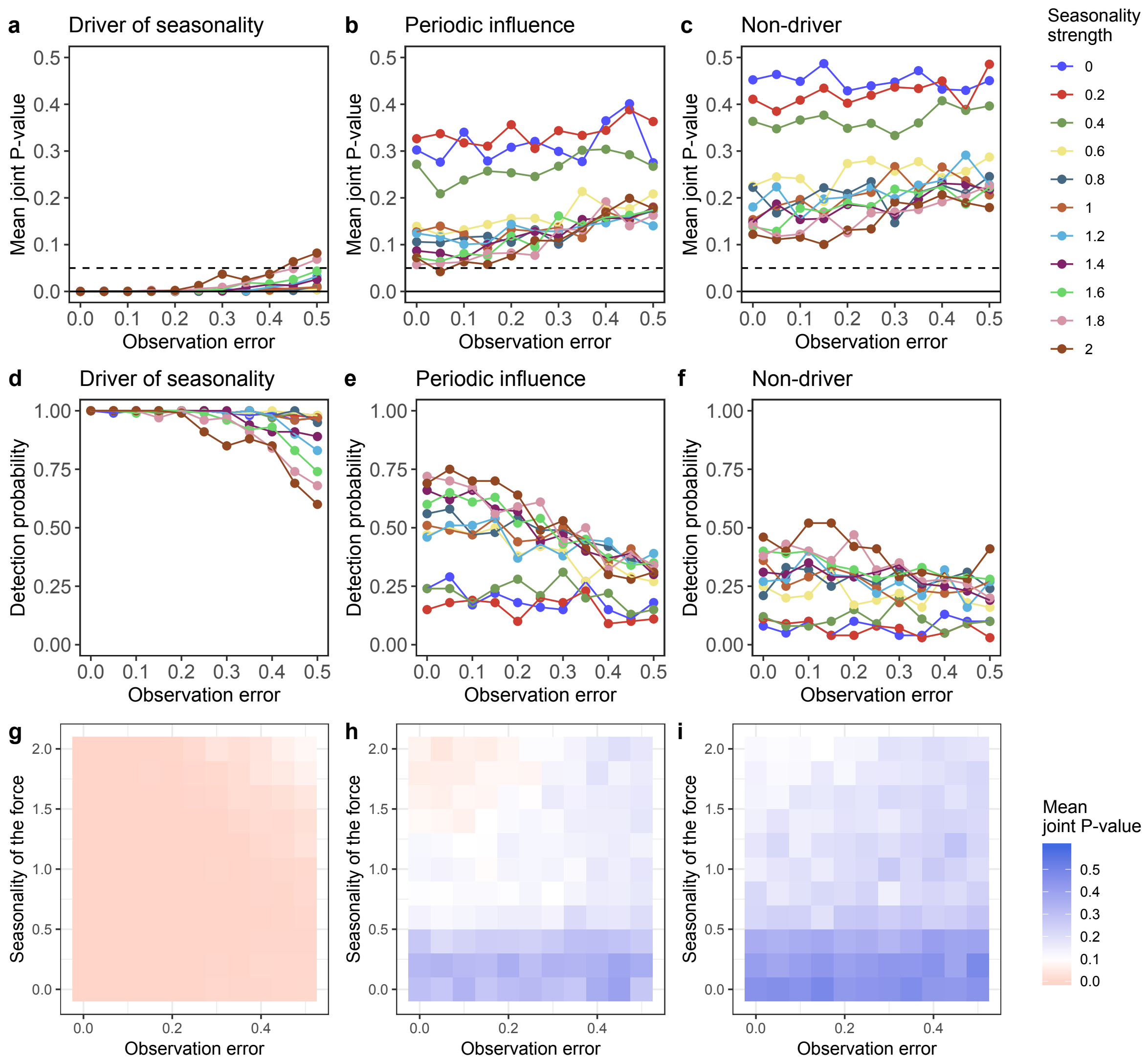

**Figure S3 | The results of the performance test of convergent cross mapping (normal CCM).** Seasonality strength, observation errors and mean joint *P*-values of normal CCM applied to time series that have (**a**) a continuous influence (the true seasonality driver), (**b**) a periodic influence, and (**c**) no influence on the target variable. Different colours indicate different strengths of seasonality. Dashed lines indicate joint *P*-value = 0.05. Seasonality strength, observation errors and detection probability (as the true seasonality driver) of normal CCM applied to time series that have (**d**) a continuous influence (the true seasonality driver), (**e**) a periodic influence, and (**f**) no influence on the target variable. Different colours indicate different strengths of seasonality. Heat map of the seasonality strength, observation errors and mean joint P-values of normal CCM applied to time series that have (**g**) a continuous influence (the true seasonality driver), (**h**) a periodic influence, and (**i**) no influence on the target variable. Colours indicate mean joint *P*-value. If time series has weak to strong seasonality, then normal CCM may incorrectly detect a non-driver as the driver (**b**–**c**). Also, even with weak to moderate strength of seasonality, the probability of false-positive detection is high for time series that have periodic influence (50–75%; **e**) and no influence (20–50%; **f**).

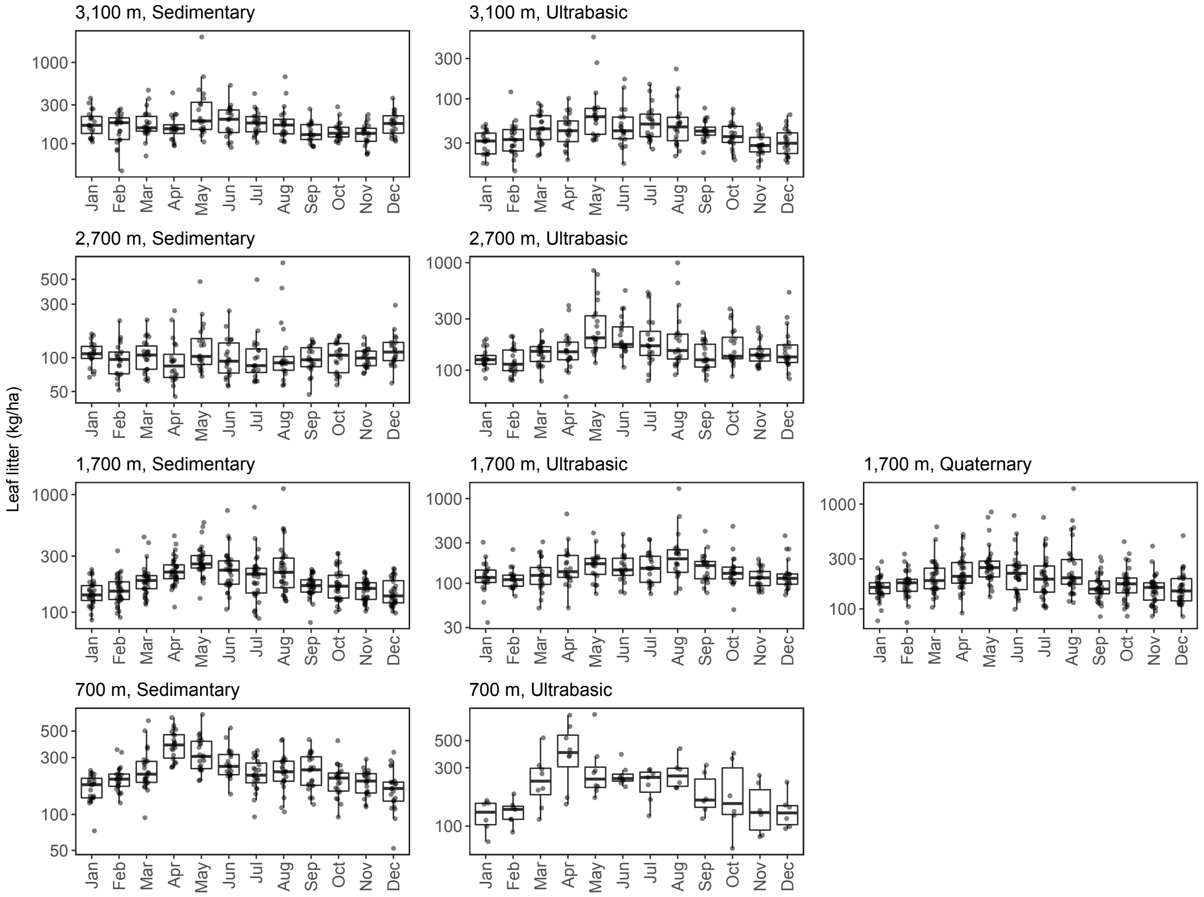

**Figure S4 | Monthly patterns of leaf litter production in the tropical rain forests on Mt. Kinabalu.** In general, leaf litter production is larger from March to June than in the other months at most sites. This trend is detected as the one-year periodicity using Fourier analysis.

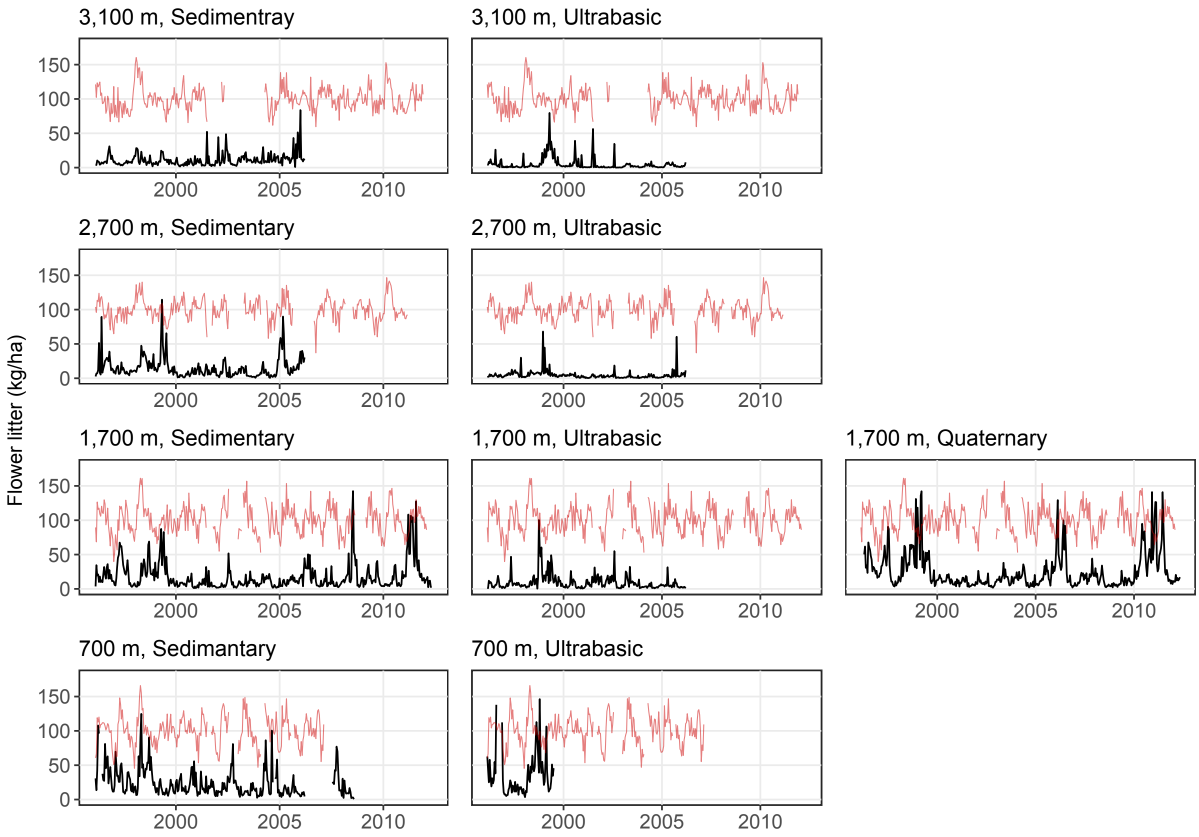

**Figure S5 | Time series of flower litter production in tropical rain forests on Mt. Kinabalu.** Black lines indicate flower litter production, and red lines indicate mean daily air temperature corrected by a general additive model (only patterns are shown). The values on the y-axis are for flower litter production. Flower litter production does not show clear annual periodicity, unlike leaf litter production.

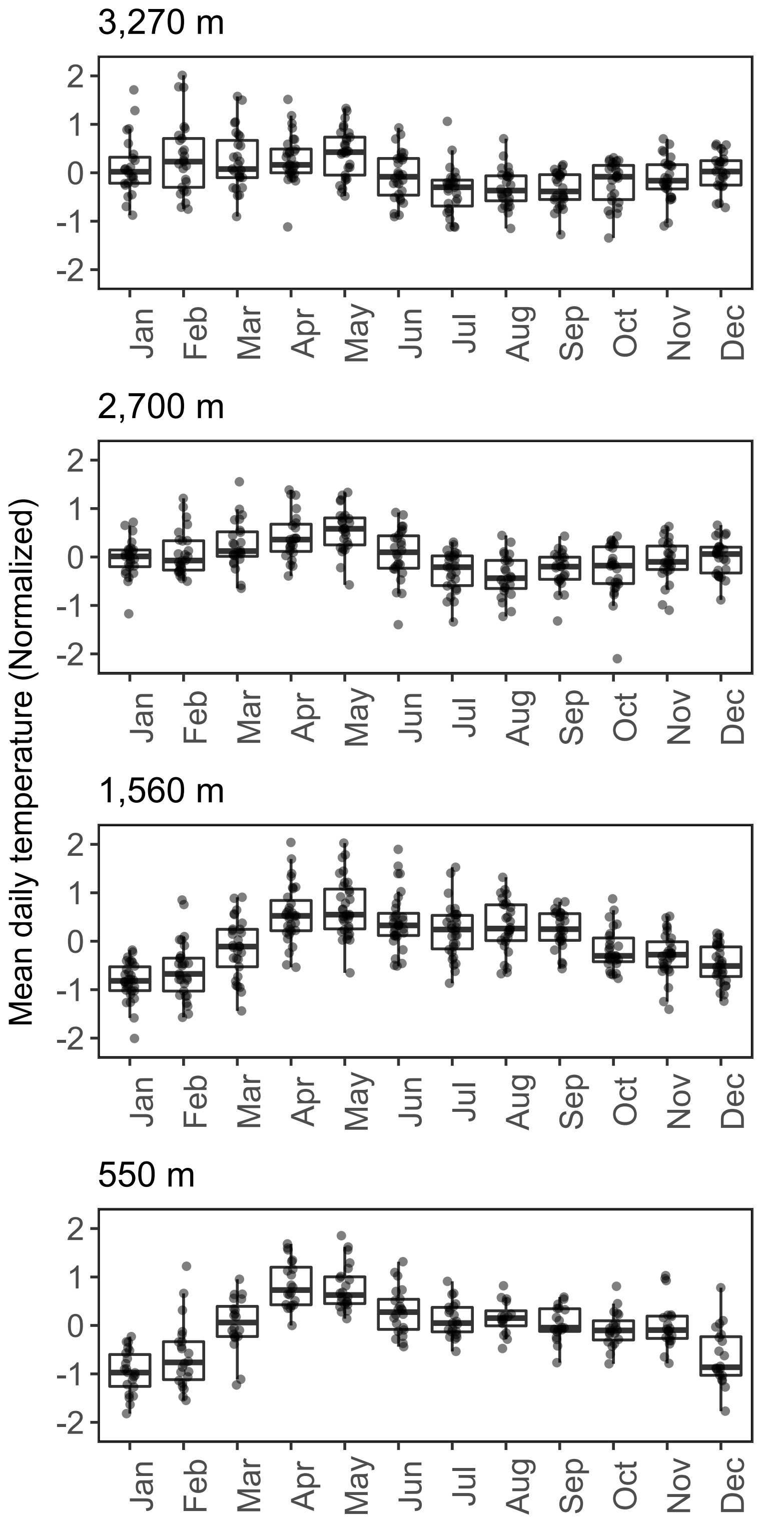

**Figure S6 | Monthly patterns of daily mean air temperature.** The original values were corrected by a general additive model and the residuals are shown. In general, daily mean air temperature is higher from March to June.

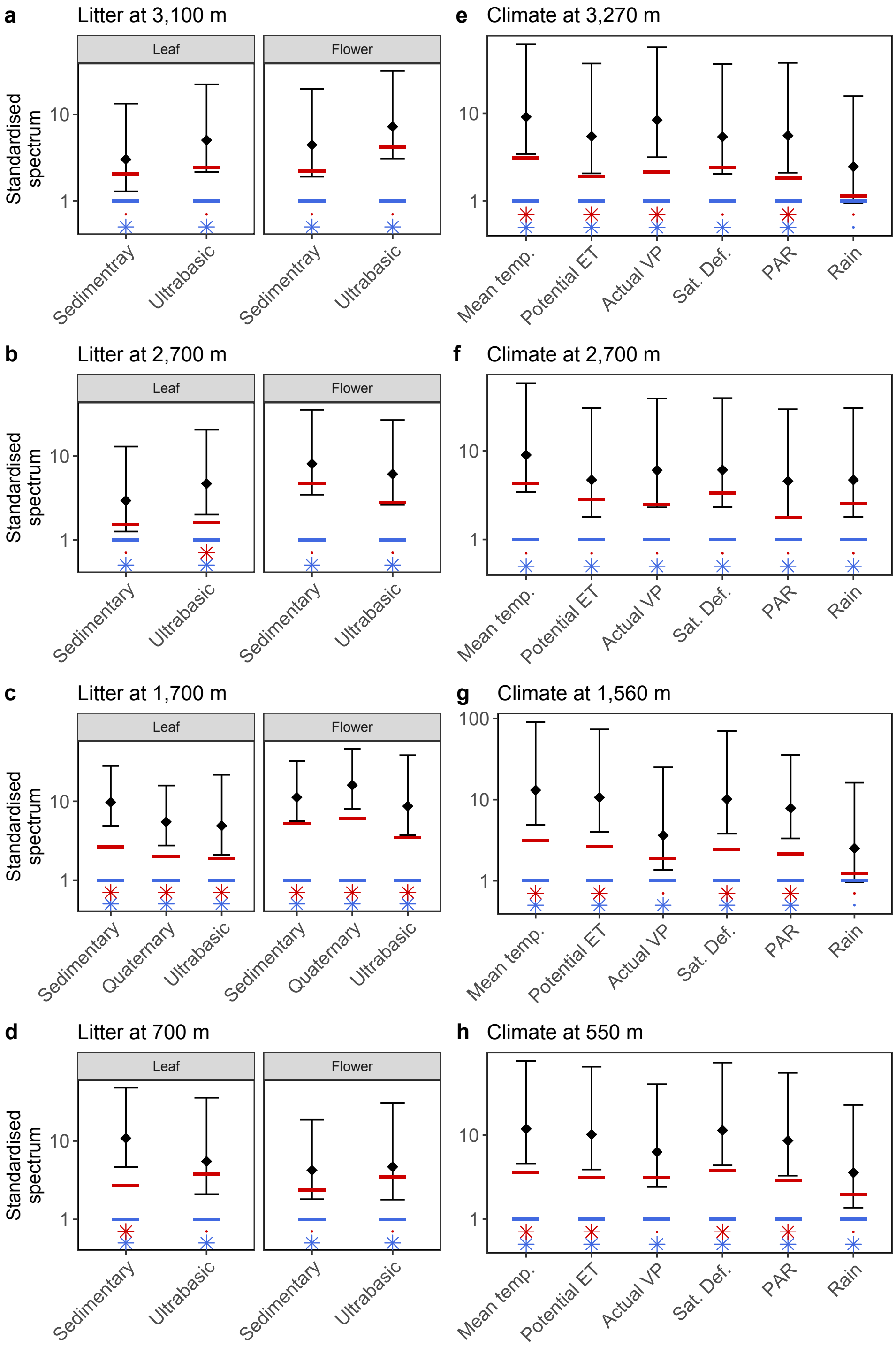

**Figure S7 | Confidence intervals and significance of the dominant periodicity of litter and meteorological time series.** Black points indicate the power of the dominant periodicity (note that the powers were not always for the 1-year periodicity). Bars indicate 95% confidence intervals calculated by assuming that spectral estimates approximate a chi-square distribution. Red and blue bars indicate null and average spectrum, respectively. Red and blue asterisks indicate the significance of the power based on the null and average spectrum, respectively. If the power is not significant, only ”dot” is shown.

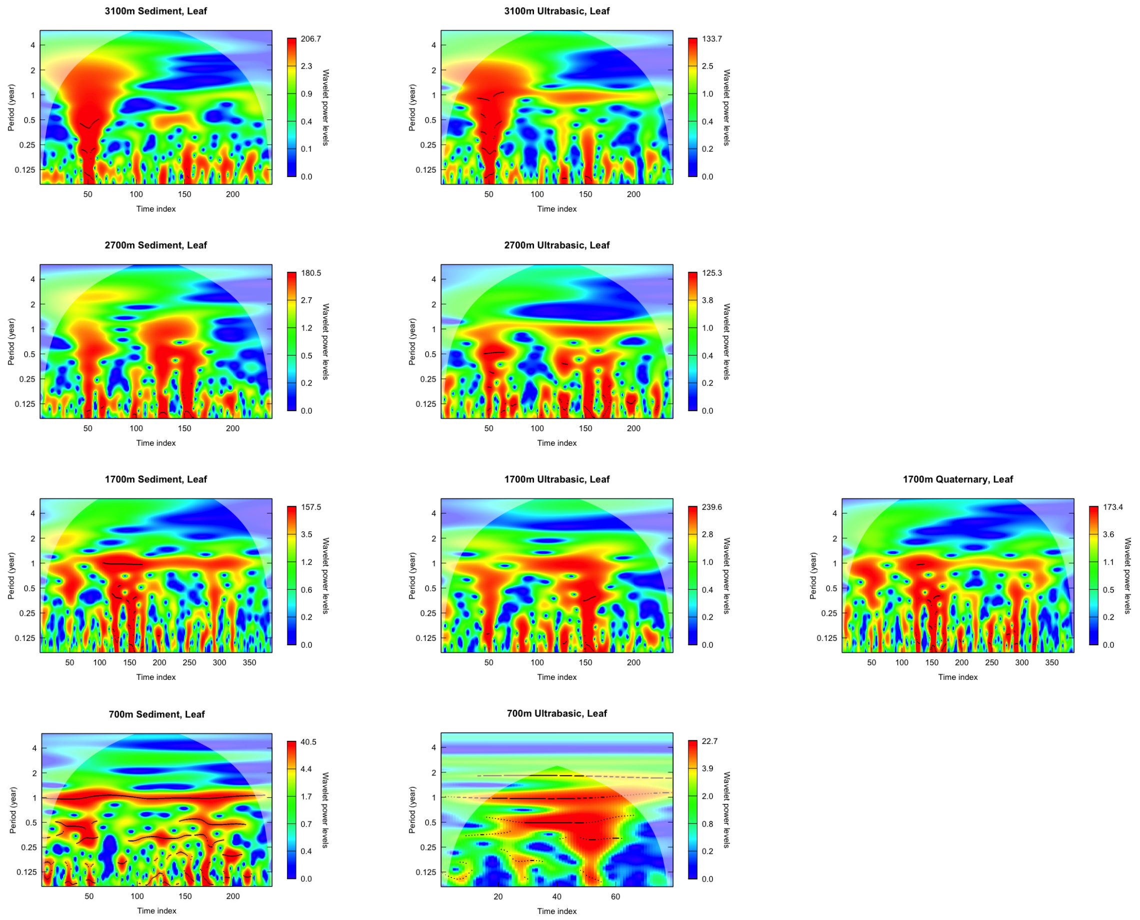

**Figure S8 | Wavelet analysis of leaf litter production.** The *x*-axis and the *y*-axis and color indicate time points, the length of periodicity and the strength of the periodicity at each frequency, respectively. Black lines indicate the ridge of wavelet power.

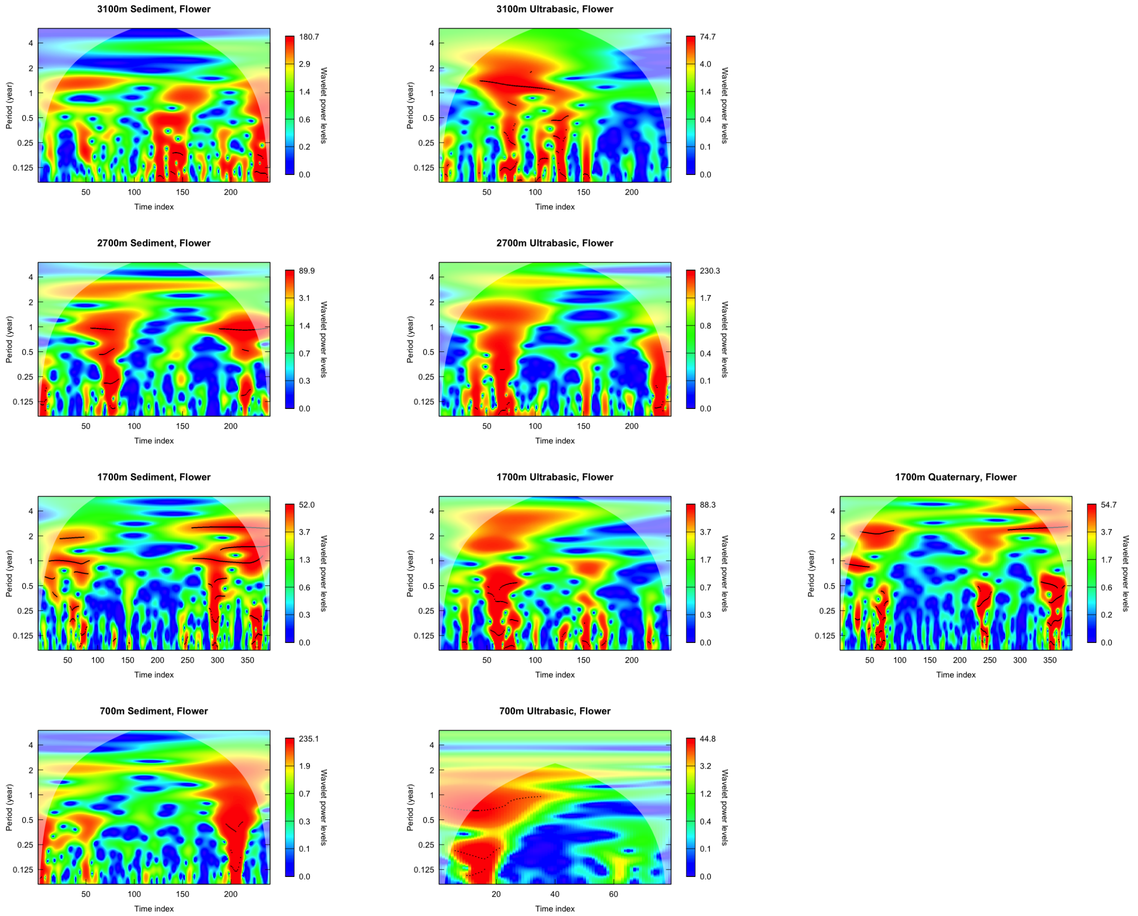

**Figure S9 | Wavelet analysis of flower litter production.** The *x*-axis and the *y*-axis and color indicate time points, the length of periodicity and the strength of the periodicity at each frequency, respectively. Black lines indicate the ridge of wavelet power.

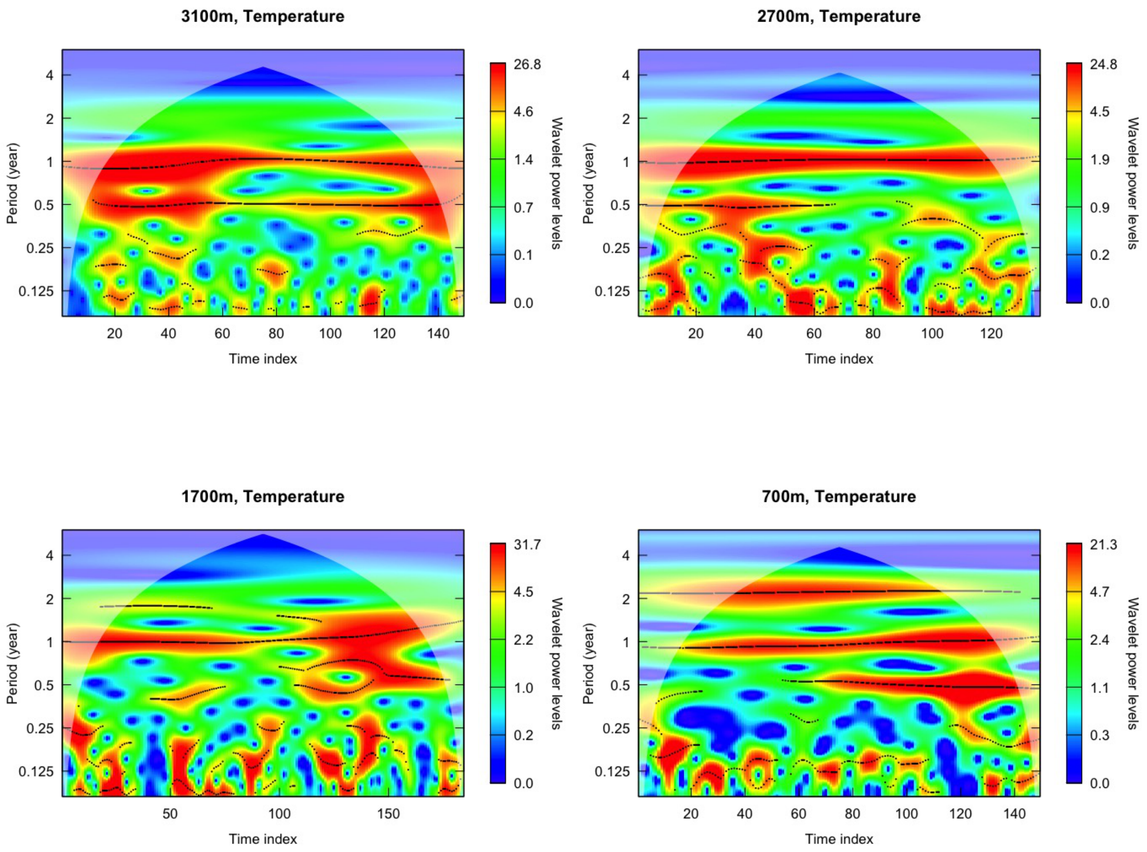

**Figure S10 | Wavelet analysis of daily mean air temperature.** The $x$-axis and the $y$-axis and color indicate time points, the length of periodicity and the strength of the periodicity at each frequency, respectively. Black lines indicate the ridge of wavelet power.

**Table S1 | Time series length used for Fourier analysis**

| **Altitude (m)** | **Parent materia** | **Total monitoring duratio** | **For Fourier analysis Start (year/month)** | **End (year/month)** | **duration (mo)** | **Time poin (*N*)** |
|---|---|---|---|---|---|---|
| *Climate site* | | | (for air temperature*) | | | |
| 3270 | | 1996/2 – 2011/12 | 2004/4 | 2011/12 | 92.0 | 184 |
| 2700 | | 1996/2 – 2011/3 | 1996/5 | 2002/7 | 74.5 | 149 |
| 1560 | | 1996/2 – 2012/2 | 2003/1 | 2008/8 | 68.0 | 136 |
| 550 | | 1996/2 – 2007/2 | 1996/5 | 2002/7 | 74.5 | 149 |
| *Litter site* | | | (for leaf and flower litter) | | | |
| 700 | Sedimentary | 1996/4 – 2006/3 | 1996/4 | 2006/3 | 120.0 | 240 |
| 1700 | Sedimentary | 1996/4 – 2012/5 | 1996/4 | 2012/5 | 193.5 | 387 |
| 2700 | Sedimentary | 1996/4 – 2006/3 | 1996/4 | 2006/3 | 120.0 | 240 |
| 3100 | Sedimentary | 1996/4 – 2006/3 | 1996/4 | 2006/3 | 120.0 | 240 |
| 700 | Ultrabasic | 1996/4 – 1999/7 | 1996/4 | 1999/7 | 39.5 | 79 |
| 1700 | Ultrabasic | 1996/4 – 2006/3 | 1996/4 | 2006/3 | 120.0 | 240 |
| 2700 | Ultrabasic | 1996/4 – 2006/3 | 1996/4 | 2006/3 | 120.0 | 240 |
| 3100 | Ultrabasic | 1996/4 – 2006/3 | 1996/4 | 2006/3 | 120.0 | 240 |
| 1700 | Quaternary | 1996/4 – 2012/5 | 1996/4 | 2012/5 | 193.0 | 386 |

*The duration may slightly change depending on meteorological variables.

**Table S2 | Summary of Fourier analysis of litter**

| Site name | Altitude (m) | Parent material | Variable | Dominant cycle (month) | Significance of the cycl (null) | (average) | One-year cycle† (12 ± 1.5 month) |
|---|---|---|---|---|---|---|---|
| POR | 700 | Sedimantary | Leaf | 12.0 | * | * | One-year cycle |
| NAL | 700 | Ultramafic | Leaf | 13.3 | N.S. | * | 7 / 9 sites |
| PHQ | 1700 | Sedimantary | Leaf | 11.8 | * | * | |
| BAB | 1700 | Ultramafic | Leaf | 12.0 | * | * | Significant one-year cycle |
| ULA | 1700 | Quaternary | Leaf | 12.0 | * | * | (null spectrum) 5 / 9 sites |
| RTM | 2700 | Sedimantary | Leaf | 6.3 | N.S. | * | (avr. spectrum) 7 / 9 sites |
| CAR | 2700 | Ultramafic | Leaf | 12.0 | N.S. | * | |
| PAK | 3100 | Sedimantary | Leaf | 24.0 | N.S. | * | |
| HEL | 3100 | Ultramafic | Leaf | 11.8 | * | * | |
| POR | 700 | Sedimantary | Flower | 24.0 | N.S. | * | One-year cycle |
| NAL | 700 | Ultramafic | Flower | 20.0 | N.S. | * | 0 / 9 sites |
| PHQ | 1700 | Sedimantary | Flower | >48 | * | * | |
| BAB | 1700 | Ultramafic | Flower | 40.0 | * | * | Significant one-year cycle |
| ULA | 1700 | Quaternary | Flower | >48 | N.S. | * | (null spectrum) 0 / 9 sites |
| RTM | 2700 | Sedimantary | Flower | 40.0 | N.S. | * | (avr. spectrum) 0 / 9 sites |
| CAR | 2700 | Ultramafic | Flower | 40.0 | N.S. | * | |
| PAK | 3100 | Sedimantary | Flower | 15.0 | N.S. | * | |
| HEL | 3100 | Ultramafic | Flower | >48 | * | * | |
| POR | 700 | Sedimantary | Branch | 4.0 | N.S. | * | One-year cycle |
| NAL | 700 | Ultramafic | Branch | 1.9 | N.S. | * | 4 / 9 sites |
| PHQ | 1700 | Sedimantary | Branch | 11.8 | * | * | |
| BAB | 1700 | Ultramafic | Branch | >48 | N.S. | * | Significant one-year cycle |
| ULA | 1700 | Quaternary | Branch | 12.0 | * | * | (null spectrum) 3 / 9 sites |
| RTM | 2700 | Sedimantary | Branch | 12.0 | N.S. | * | (avr. spectrum) 4 / 9 sites |
| CAR | 2700 | Ultramafic | Branch | 12.0 | * | * | |
| PAK | 3100 | Sedimantary | Branch | 6.0 | * | * | |
| HEL | 3100 | Ultramafic | Branch | 1.6 | N.S. | N.S. | |
| POR | 700 | Sedimantary | Epiphyte | 40.0 | N.S. | * | One-year cycle |
| NAL | 700 | Ultramafic | Epiphyte | 13.3 | N.S. | N.S. | 3 / 9 sites |
| PHQ | 1700 | Sedimantary | Epiphyte | 12.5 | N.S. | * | |
| BAB | 1700 | Ultramafic | Epiphyte | 30.0 | N.S. | * | Significant one-year cycle |
| ULA | 1700 | Quaternary | Epiphyte | 12.0 | * | * | (null spectrum) 1 / 9 sites |
| RTM | 2700 | Sedimantary | Epiphyte | 1.0 | N.S. | * | (avr. spectrum) 2 / 9 sites |
| CAR | 2700 | Ultramafic | Epiphyte | >48 | N.S. | N.S. | |
| PAK | 3100 | Sedimantary | Epiphyte | 17.1 | N.S. | * | |
| HEL | 3100 | Ultramafic | Epiphyte | 10.0 | N.S. | * | |
| POR | 700 | Sedimantary | Bamboo | 1.1 | * | * | One-year cycle |
| NAL | 700 | Ultramafic | Bamboo | 13.3 | N.S. | * | 2 / 9 sites |
| PHQ | 1700 | Sedimantary | Bamboo | >48 | * | * | |
| BAB | 1700 | Ultramafic | Bamboo | 1.0 | N.S. | N.S. | Significant one-year cycle |
| ULA | 1700 | Quaternary | Bamboo | 12.0 | N.S. | * | (null spectrum) 0 / 9 sites |
| RTM | 2700 | Sedimantary | Bamboo | >48 | N.S. | * | (avr. spectrum) 1 / 9 sites |
| CAR | 2700 | Ultramafic | Bamboo | >48 | N.S. | N.S. | |
| PAK | 3100 | Sedimantary | Bamboo | 9.2 | N.S. | N.S. | |
| HEL | 3100 | Ultramafic | Bamboo | >48 | * | * | |
| POR | 700 | Sedimantary | Dust | >48 | N.S. | * | One-year cycle |
| NAL | 700 | Ultramafic | Dust | 40.0 | N.S. | * | 4 / 9 sites |
| PHQ | 1700 | Sedimantary | Dust | 11.8 | * | * | |
| BAB | 1700 | Ultramafic | Dust | >48 | N.S. | * | Significant one-year cycle |
| ULA | 1700 | Quaternary | Dust | 12.0 | * | * | (null spectrum) 4 / 9 sites |
| RTM | 2700 | Sedimantary | Dust | 12.0 | * | * | (avr. spectrum) 4 / 9 sites |
| CAR | 2700 | Ultramafic | Dust | >48 | * | * | |
| PAK | 3100 | Sedimantary | Dust | >48 | * | * | |
| HEL | 3100 | Ultramafic | Dust | 11.8 | * | * | |

†This column indicates how many study sites exhibited annual seasonality (i.e., one-year cycle). "One-year cycle" indicates the number of study sites of which dominant cycle is 12 ± 1.5 months. "Significant one-year cycle" indicates the number of study sites which showed significant annual seasonality compared with null or average spectrum.

**Table S3 | Summary of Fourier analysis of climate variables**

| Site name | Altitude (m) | Variable | Dominant cycle (month) | Significance of the cycl (null) | (average) | One-year cycle† (12 ± 1.5 month) |
|---|---|---|---|---|---|---|
| POR | 550 | Mean daily temperatu | 12.5 | * | * | One-year cycle |
| POR | 550 | Relative humidity | 12.5 | * | * | 5 / 7 variables |
| POR | 550 | Acutual vapor pressur | 15.0 | N.S. | * | |
| POR | 550 | Saturation deficient | 12.5 | * | * | Significant one-year cycle |
| POR | 550 | PAR | 12.5 | * | * | (null spectrum) 5 / 7 variables |
| POR | 550 | Evapotranspiration | 12.5 | * | * | (avr. spectrum) 7 / 7 variables |
| POR | 550 | Rain | 7.5 | N.S. | * | |
| PHQ | 1650 | Mean daily temperatu | 12.0 | * | * | One-year cycle |
| PHQ | 1650 | Relative humidity | 12.0 | * | * | 4 / 7 variables |
| PHQ | 1650 | Acutual vapor pressur | 6.0 | * | * | |
| PHQ | 1650 | Saturation deficient | 12.0 | N.S. | * | Significant one-year cycle |
| PHQ | 1650 | PAR | 5.9 | * | * | (null spectrum) 3 / 7 variables |
| PHQ | 1650 | Evapotranspiration | 12.0 | * | * | (avr. spectrum) 4 / 7 variables |
| PHQ | 1650 | Rain | 1.7 | N.S. | N.S. | |
| CAR | 2700 | Mean daily temperatu | 25.0 | N.S. | * | One-year cycle |
| CAR | 2700 | Relative humidity | 12.5 | N.S. | * | 2 / 7 variables |
| CAR | 2700 | Acutual vapor pressur | 6.3 | N.S. | * | |
| CAR | 2700 | Saturation deficient | 12.5 | N.S. | * | Significant one-year cycle |
| CAR | 2700 | PAR | 6.3 | N.S. | * | (null spectrum) 0 / 7 variables |
| CAR | 2700 | Evapotranspiration | 8.4 | N.S. | * | (avr. spectrum) 2 / 7 variables |
| CAR | 2700 | Rain | 25.0 | N.S. | * | |
| LAB | 3270 | Mean daily temperatu | 12.0 | * | * | One-year cycle |
| LAB | 3270 | Relative humidity | 12.0 | N.S. | * | 3 / 7 variables |
| LAB | 3270 | Acutual vapor pressur | 6.0 | * | * | |
| LAB | 3270 | Saturation deficient | 12.0 | N.S. | * | Significant one-year cycle |
| LAB | 3270 | PAR | 6.0 | * | * | (null spectrum) 1 / 7 variables |
| LAB | 3270 | Evapotranspiration | 6.0 | * | * | (avr. spectrum) 3 / 7 variables |
| LAB | 3270 | Rain | 2.7 | N.S. | N.S. | |

†This column indicates how many study sites exhibited annual seasonality (i.e., one-year cycle). "One-year cycle" indicates the number of study sites of which dominant cycle is 12 ± 1.5 months. "Significant one-year cycle" indicates the number of study sites which showed significant annual seasonality compared with null or average spectrum.
